# Drone-based effective counting and ageing of hippopotamus (*Hippopotamus amphibius*) in the Okavango Delta in Botswana

**DOI:** 10.1101/689059

**Authors:** Victoria L. Inman, Richard T. Kingsford, Michael J. Chase, Keith E. A. Leggett

## Abstract

Accurately estimating hippopotamus (*Hippopotamus amphibius*) numbers is difficult due to their aggressive nature, amphibious lifestyle, and habit of diving and surfacing. Traditionally, hippos are counted using aerial surveys and land/boat surveys. We compared estimates of numbers of hippos in a lagoon in the Okavango Delta, counted from land and video taken from a DJI Phantom 4™ drone, testing for effectiveness at three heights (40 m, 80 m, and 120 m) and four times of day (early morning, late morning, early afternoon, and late afternoon). In addition, we determined effectiveness for differentiating age classes (juvenile, subadult, and adult), based on visual assessment and measurements from drone images, at different times and heights. Estimates in the pool averaged 9.18 (± 0.25SE, range 1 – 14, n = 112 counts). Drone counts at 40 m produced the highest counts of hippos, 10.6% higher than land counts and drone counts at 80 m, and 17.6% higher than drone counts at 120 m. Fewer hippos were counted in the early morning, when the hippos were active and most likely submerged, compared to all other times of day, when they tended to rest in shallow water with their bodies exposed. We were able to assign age classes to similar numbers of hippos from land counts and counts at 40 m, although land counts were better at identifying juveniles and subadults. Early morning was the least effective time to age hippos given their active behaviour, increasingly problematic with increasing height. Use of a relatively low-cost drone provided a rigorous and repeatable method for estimating numbers and ages of hippos, but not in the early morning.

## Introduction

Hippopotamus or hippo (*Hippopotamus amphibius*) are Vulnerable on the IUCN Red List of Threatened Species, numbering 115,000–130,000 [1]. Habitat loss and hunting for meat and ivory are driving declines [1], but much of the population data originates from aerial surveys, which can be inaccurate [2–6]. Reliable and accurate spatial and temporal data on abundances and demographics of hippo populations are essential for effective conservation management [1,7,8] but hippos are inherently difficult to count because individuals regularly submerge and surface throughout the day and have uniform appearance. They are also among the more dangerous animals in Africa [9–11], limiting effectiveness of on-land and water methods of counting [12].

Hippos are usually surveyed from the air [9–12], but also from boats and land [9,13,14]; each method has advantages and disadvantages. Aerial surveys cover large areas [4] but with limited time to scan waterbodies and count hippos, given their speed. Also, aircraft noise may cause hippos to submerge [19], contributing to underestimation [20]. Aerial surveys are costly and logistically difficult, resulting in long intervals between surveys [21–24]. Slow, low-flying microlight aircraft or helicopters capturing images may overcome some of these challenges [25] but remain costly and potentially logistically difficult, often still causing disturbance. Counts from land or boats tend to be more accurate [4], given hippo pods can be observed for a long period of time, allowing for differential submergence, but such counts can be dangerous and difficult or impossible to do where hippo pods are in remote or not easily accessible areas [21,26]. Further, line of sight from horizontal counts can be challenging, given some individuals are inevitably obscured by others [27], even though accuracy improves when hippos rest in aggregations, where most of their bodies are visible (“rafting”) [28]. Such difficulties compound when assessing demographic composition of hippo pods.

Drones (unmanned aerial systems/vehicles or remotely piloted aircraft) are an increasingly effective means for monitoring animals, including birds [29], turtles [30], dugongs [23], and cetaceans [31]. They usually have low impact, are relatively low cost, have consistent flight paths, allow remote operation away from wildlife, and enable monitoring of areas inaccessible by land or boat [23,30,32,33]. Hippos were counted, including their demographic composition, in pods in the Democratic Republic of Congo, using relatively expensive technology and sophisticated methods [34–36], but without comparing drone counts to a current survey method. Drone height and weather affected hippo detection, based on surveys only done in the early morning [36]. However, time of day is critical, given hippo behaviour varies throughout the day [3,20,37–40]. We trialled the use of a relatively low-cost drone, testing its effectiveness to estimate hippo numbers, the percentage of hippos that could be assigned to age classes, and numbers of juveniles, subadults, and adults, comparing these estimates to counts from land. We also tested how time of day and survey height affected these counts.

## METHODS

We flew a drone (1380 gram multirotor DJI Phantom 4™ (www.dji.com), 4K-quality video, 12.4 MP photo, aperture of f/2.8) over a lagoon (−19.41725°E, 22.56815°S, 2.4 ha), with a resident hippo population, within NG26 (Abu Concession) of the Okavango Delta, northern Botswana (Fig. 1), on 7^th^, 8^th^, 11^th^, 13^th^ and 14^th^ November 2017 and 2^nd^ and 3^rd^ December 2017. The camera was controlled and stabilised by a three-axis gimbal, and the drone controlled by a GPS-stabilised system. All videos were filmed at 3840 × 2160 pixels (30 frames s^−1^), with automatic ISO and shutter speed, allowing variation for neutrally exposed images. Sensor width was 6.2 mm and camera focal length was 3.61 mm [41]. The drone was programmed to fly (5.4 km hr^−1^) in transects over the lagoon, calculated and controlled using the Drone Harmony app (www.droneharmony.com) and run through a smartphone, while continuously recording video. The lagoon was outlined using the satellite imagery provided on the app, and routes automatically calculated to ensure the entire surface area of the lagoon was captured on video, with a horizontal overlap of 20 percent, with the camera facing directly downward (gimbal angle of −90°).

**Fig 1.**
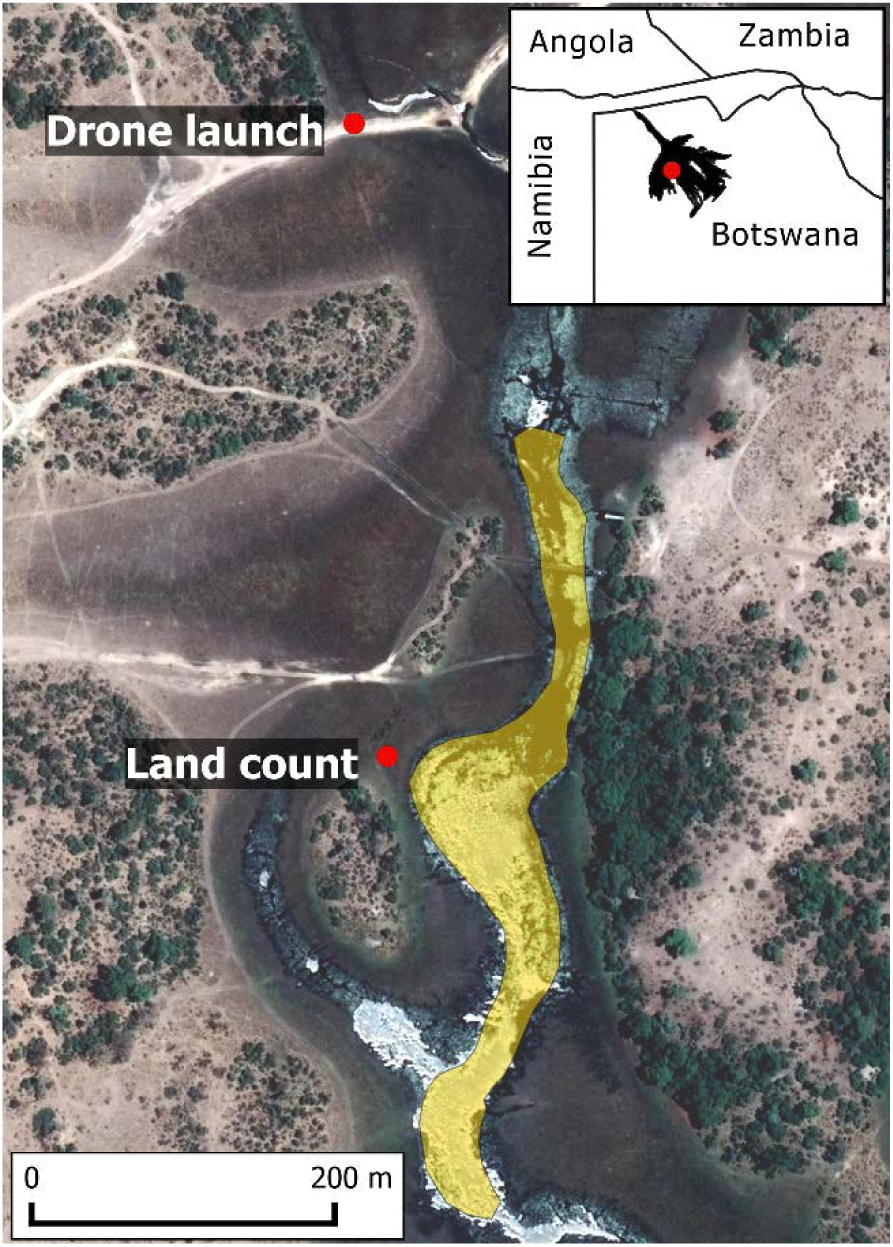
Lagoon (shaded yellow) where hippos were counted in NG26, within the Okavango Delta in Botswana, showing where the drone was launched and land counts were done.

## DRONE SURVEYS

We flew the drone at three heights sequentially, in descending order (120 m, 80 m, 40 m), starting with the height least likely to cause disturbance. The drone was launched and landed out of visual range of the hippos (Fig. 1), avoiding disturbing them, only returning after two flights to change the battery. It took 30-40 minutes to complete the drone surveys. Given differential coverage of the lagoon, routes varied from backwards and forwards (east-west) across the lagoon (40 m, 80 m) to one path down the centre, north-south (120 m). Surveys were conducted four times a day (early morning 6:30 – 7:30 [EM]; late morning 10:00-11:00 [LM]; early afternoon 13:30 – 14:30 [EA]; late afternoon 17:00 – 18:00 [LA]), evenly dividing diurnal hours from an hour after sunrise to an hour before sunset, when there was maximum visibility. We completed a total of 84 drone counts (28 per height, 21 per day). For estimating numbers and measuring hippos, all 84 flight videos were randomised to avoid bias among proximate assessments. When reviewing the videos, we also checked for behaviours indicating the hippos were disturbed by the drone.

## LAND SURVEYS

We also counted hippos from a vehicle on land adjacent to the lagoon with two people (15 mins), in the same location each time (Fig. 1) where all hippos could be observed, immediately following the last drone flight for each time of day (i.e. four land counts a day). We noted any behaviours indicating hippos were disturbed by our presence, including submerging in water, vocalising, yawning, and charging [12,42]. We completed a total of 28 land counts (seven per day).

### Measurement for ageing

Ground sampling distance (GSD, e.g. pixel size) was calculated based on the equation:

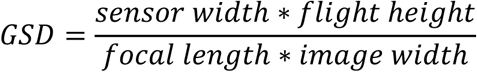

We used the ‘snapshot’ function of VLC media player [43] to obtain still images of each hippo visible on video. Individual images were imported into ImageJ [44], the ‘set scale’ function used to input the GSD for that image (1.79cm/pixel for drone images at 40 m, 3.58cm/pixel at 80 m, and 5.37cm/pixel at 120 m) and the ‘straight line’ function used to measure the length of each hippo from the tip of the snout to the base of the tail. This measured length was then used to assign each hippo to an age class, with no differentiation between males and females, based on the known relationship between body length and age [45]. Hippos < 184 cm less than two years old were classed as juveniles (hippos produce a calf about once every two years [46]); hippos 184 – 233 cm long were two to four years old and classed as subadults, and hippos > 233 cm were classed as adults, given lower-end estimates of age of puberty in hippos is four years [45,47–49]. If the entire body was not visible (e.g. the hippo was partially submerged), but the visible section exceeded 233 cm, then it was classed as an adult. Other partially submerged hippos, where the snout and base of the tail were not visible, were classed ‘unknown’. For land counts, hippos were similarly assigned into the three age classes. Hippos judged as less than 1/2 the length of the largest hippo (typically the dominant male [50,51]) were classed as juveniles; subadults were between 1/2 and 2/3 the length of the largest hippo; and adults were over 2/3 the length of the largest hippo. Based on the proposed maximum hippo body length of 359cm [45], the land count classes aligned well with the drone classes, with the distinction between juveniles and subadults calculated as 179.50 cm (compared to 184 cm) and between subadults and adults as 239.30 cm (compared to 233 cm). During visual assessments, if hippos could not be assigned to an age class, they were recorded as unknown.

### Analysis

We tested the effect of height, time of day, and their interaction on variations in total hippo count (model 1), percentage of hippos assigned to age classes (number of juveniles, subadults, and adults divided by the total count for each drone/land survey; model 2), and counts of juveniles, subadults, and adults (models 3 - 5). ‘Height’ had four levels (land count and drone heights 40 m, 80 m, 120 m) as did ‘time of day’ (early morning [EM], late morning [LM], early afternoon [EA], and late afternoon [LA]). Height and time of day were defined as fixed effects, with survey date as a random effect. For model 1, we used a linear mixed model (lmer and anova functions, lme4 package [52]), for model 2 a generalized linear mixed-effect model (glmer function, lme4 package; Anova, car package [53]), with family Binomial and weights equal to the total number of hippos for each count, and for models 3 – 5 generalized linear models (manyglm and anova.manyglm functions, mvabund package [54]), with family Negative Binomial. Differences among the levels of the effects were tested using post hoc pairwise comparisons, based on estimated marginal means, using a Tukey adjustment with the emmeans package [55] (model 1 and 2) and the pairwise.comp function (manyglm package), adjusted for multiple testing using a step-down resampling procedure (models 3 -5). We determined the maximum number of hippos seen from any count (drone or land) for each day, as a proxy of potential true number of hippos, given that hippos generally do not move out of lagoons during diurnal hours [56], investigating how counts compared to this daily maximum, as a measure of true accuracy.

For all models, we examined plots of distributions of residuals against the predictors and Q–Q plots of the normal distribution to test the assumptions of homogeneity of variance and normality of data. These assumptions were met, requiring no transformation. All statistics were conducted using the R computing environment (version 3.5.2) [57]. Means were reported with standard errors.

### Ethics statement

This research was approved by UNSW’s Animal Care & Ethics Committee (ACEC Number 17/75A), Civil Aviation Authority of Botswana (Remotely Piloted Aircraft Certificate Number RPA (H) 147) and The Republic of Botswana Ministry of Environment, Wildlife and Tourism (Research Permit EWT 8/36/4 XXXIII (55)).

## RESULTS

The number of hippos counted in the lagoon averaged 9.18 ± 0.25 (range 1 – 14, n = 112 counts, S1 Table). The pod consisted of one juvenile, two subadults, with adults ranging in number from eleven on the first survey day to six on the last day, based on daily maximum counts. All hippos remained in the water during the surveys. The drone’s low impact sound was audible at 40 m (with decreasing noise level at higher altitudes) but hippos were not observed to be disturbed by the drone at any height, with no obvious changes in behaviour observed on the videos. The hippos were slightly disturbed by the presence of the vehicle during land counts. The hippos did not charge the vehicle or behave aggressively, but if they were near the edge of the pool when the observers arrived, they became vigilant and sometimes moved away from the observers. Their disturbance response varied with their activity, responding most when they were already active (e.g. the early morning), whereas if they were resting when we approached, they seldom moved.

Summary tables and post hoc comparisons of the fitted models are shown in Table A-E in S2 Text. There was no significant interaction between height of survey and time of day on total hippo count, so we omitted the interaction from subsequent analyses. Hippo count varied significantly with height (F_3,99_ = 4.059, *p* = 0.009) and time of day (F_3,99_ = 14.564, *p* < 0.001, Fig. 2). Hippo count was also significantly negatively related to the random effect of ‘survey date’ (χ^2^_1,9_ = 64.757, *p* < 0.001); there were 35.7% fewer hippos at the end of the survey, compared to the beginning of the surveys. Counts at 40 m were significantly higher than at 120 m (*p* = 0.005), identifying on average 17.6% more hippos (Fig. 2a). Also, 10.6% more hippos were counted at 40 m than during land counts, although this was not significant. The average number of hippos detected at 80 m was the same as the number of hippos counted from land, but numbers of hippos detected at 120 m were 5.9% less than during land counts. Early morning counts were significantly lower than at all other times of day (late morning, *p* < 0.001; early afternoon, *p* < 0.001; late afternoon, *p* < 0.001), with no significant differences among the other times of day (Fig. 2b). There were 22.4 – 26.0% fewer hippos counted during early morning counts, compared to other times of the day.

**Fig 2.**
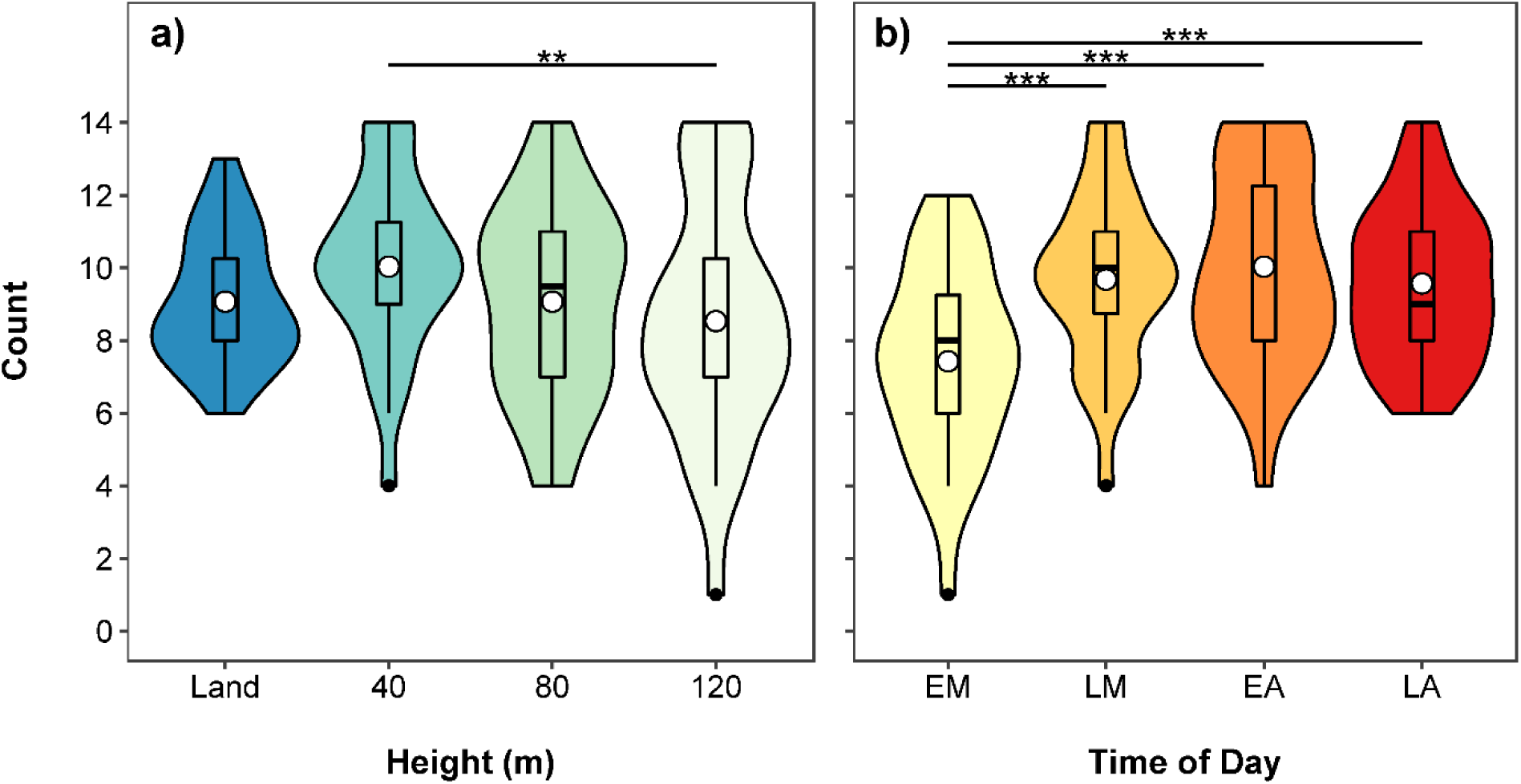
Violin and boxplot (mean, circle) showing variation in total hippo counts for a) land and three drone heights (40m, 80m and 120m) and b) time of day (EM – early morning, LM – late morning, EA – early afternoon, LA – late afternoon). Data collected using a Phantom 4 drone in a lagoon in the Okavango Delta in Botswana. Significant post hoc pairwise comparisons identified by asterisks.

Our daily maximum counts, a proxy of potential true number of hippos in the lagoon, occurred at all times of the day, although there were more in the middle of the day: early morning (3), late morning (6), early afternoon (9), and late afternoon (3, Fig. 3, Table 1). Eighteen daily maximum counts were drone counts: 40 m (10), 80 m (3), and 120 m (5), along with three land counts (Fig. 3, Table 1). The count with the greatest difference from the daily maximum was a 120 m drone count in the early morning (71.4% less hippos than daily maximum).

**Fig 3.**
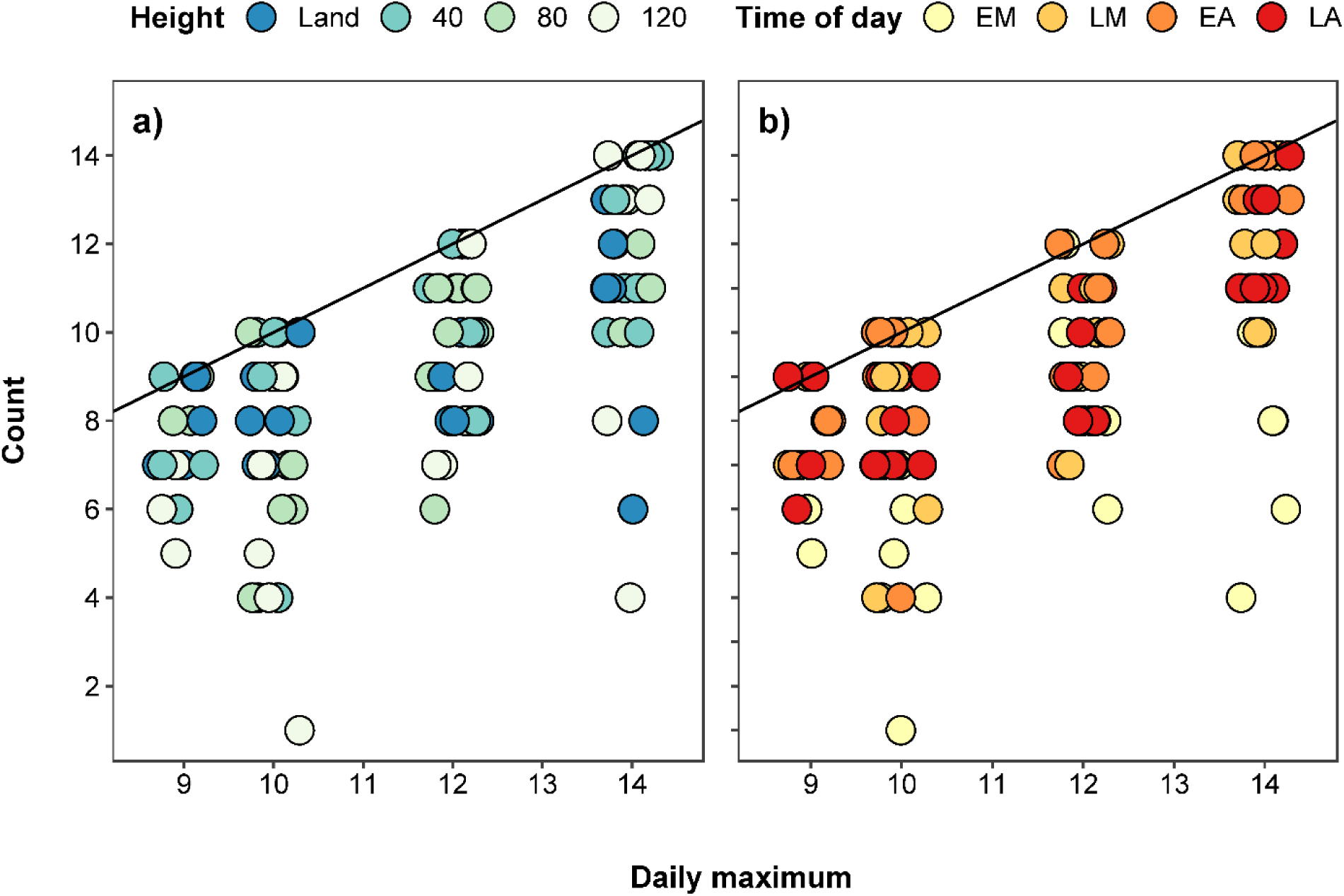
Relationship between maximum hippo counts (a proxy of the true count) and a) different heights and b) times of day (EM – early morning, LM – late morning, EA – early afternoon, LA – late afternoon). Points were jittered along the x axis.

**Table 1.** Mean counts of hippos (± SE) and number of times that each count matched the daily maximum for each observation time of day and height combination. Sample size was seven for each count.

| **Time of Day** | **Height** | **Mean count** | **Daily maximum<sup>a</sup>** |
| --- | --- | --- | --- |
| Early Morning | Land | 7.3 ± 0.3 | 1 |
| | 40 m | 9.0 ± 1.1 | 2 |
| | 80 m | 7.7 ± 1.0 | 0 |
| | 120 m | 5.7 ± 1.1 | 0 |
| Late Morning | Land | 9.7 ± 0.7 | 0 |
| | 40 m | 10.7 ± 0.7 | 4 |
| | 80 m | 8.7 ± 1.2 | 0 |
| | 120 m | 9.6 ± 0.9 | 2 |
| Early Afternoon | Land | 9.7 ± 0.7 | 1 |
| | 40 m | 10.7 ± 1.0 | 4 |
| | 80 m | 10.3 ± 1.0 | 2 |
| | 120 m | 9.4 ± 1.4 | 2 |
| Late Afternoon | Land | 9.6 ± 0.5 | 1 |
| | 40 m | 9.7 ± 0.8 | 0 |
| | 80 m | 9.6 ± 0.8 | 1 |
| | 120 m | 9.4 ± 1.2 | 1 |

<sup>a</sup> The number of times this time of day/height combination was the count with the maximum number of hippos seen for that day.

The percentage of hippos that were assigned to age classes was significantly related to height (χ^2^_3,90_ = 29.419, *p* < 0.001), time of day (χ^2^_3,90_ = 34.576, *p* < 0.001), their interaction (χ^2^_9,90_= 21.527, *p* = 0.011, Fig. 4), and survey date (χ^2^_1,17_ = 23.501, *p* < 0.001). Land counts and counts at 40 m assigned similar numbers of hippos to age classes and this did not differ with time of day. In the morning, land counts assigned more hippos to age classes than counts at 80 m (early morning, *p* < 0.001) and 120 m (early morning, *p* < 0.001; late morning, *p* = 0.012), with counts at 80 m and 120 m having significantly fewer hippos assigned to age classes in the early morning compared to all other times of day (all p < 0.05). By the early and late afternoon, all survey heights assigned similar numbers of hippos to age classes. The height and time of day survey with the highest average percentage of hippos assigned to age classes was land counts, in the late afternoon (66.8% of hippos), compared to the lowest average of 3.6% from surveys at 120 m in the early morning.

**Fig 4.**
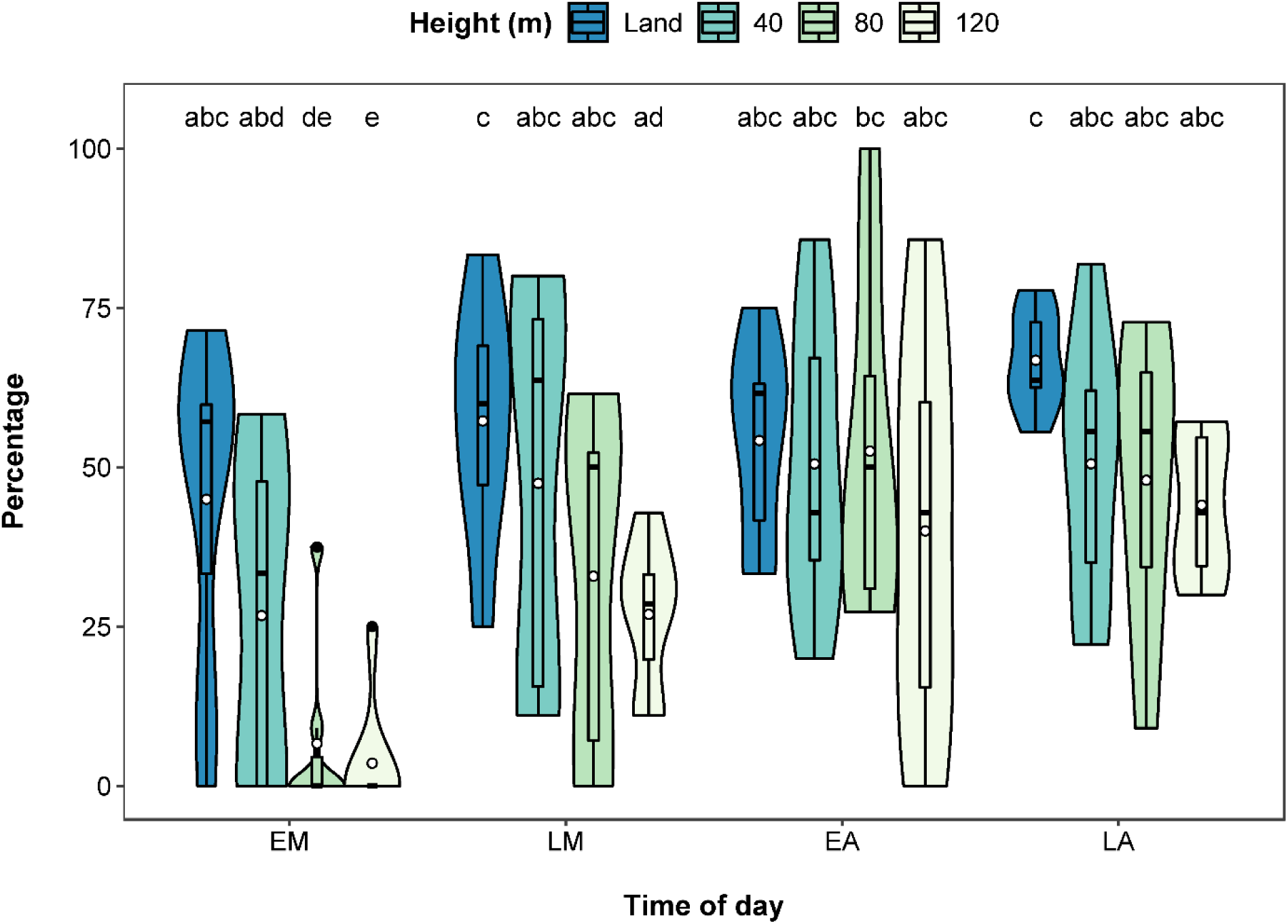
Violin and boxplot (mean, circle) showing significant interactive effect of height and time of day (EM – early morning, LM – late morning, EA – early afternoon, LA – late afternoon) on percentage of hippos assigned to the three age classes (juvenile, subadult and adult). Significant post hoc pairwise comparisons identified by letters.

There was no significant interaction between height and time of day on the number of observed juveniles or subadults and so we omitted the interaction from subsequent analyses. The number of juveniles and subadults was significantly related to height (juveniles, LRT _3,102_ = 19.173, *p* = 0.001; subadults, LRT _3,102_ = 24.151, *p* = 0.001, Fig. 5a). For both juveniles and subadults, land counts provided significantly higher counts than counts at 40 m (juveniles, *p* = 0.017; subadults, *p* = 0.008), 80 m (juveniles, *p* = 0.009; subadults, *p* = 0.029) and 120 m (juveniles, *p* = 0.001; subadults, *p* = 0.001). There were no significant differences among the other drone heights. The number of juveniles was not related to time of day but number of subadults was (LRT _3,99_ = 10.896, *p* = 0.013, Fig. 5b). Early morning counts of subadults were significantly lower than counts in the late afternoon (*p* = 0.009), with no significant differences among the other times of day. There was no effect of survey date on the number of juveniles or subadults.

**Fig 5.**
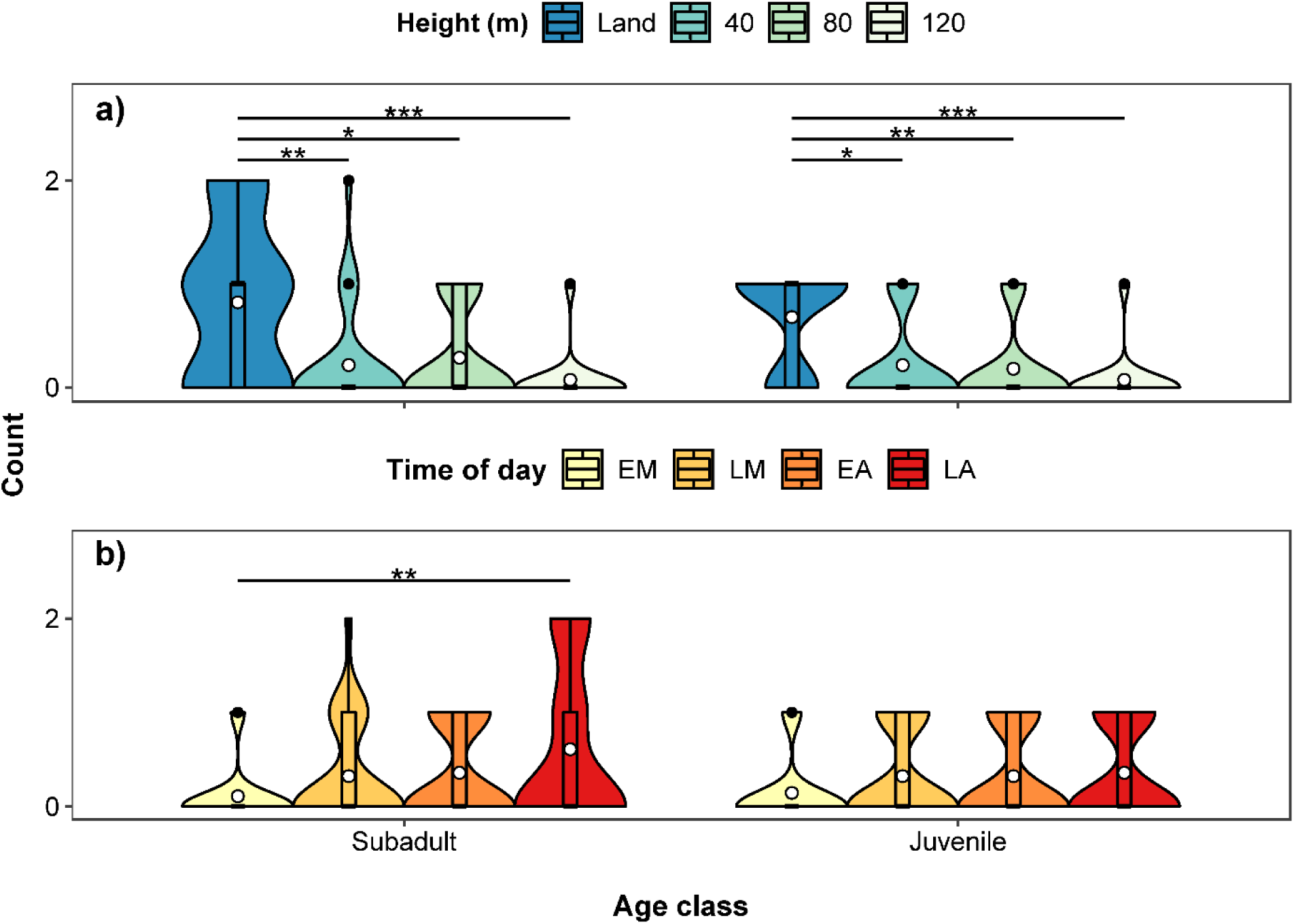
Violin and boxplot (mean, circle) showing variation in counts for juvenile and subadult hippos for a) height and b) time of day (EM – early morning, LM – late morning, EA – early afternoon, LA – late afternoon). Significant post hoc pairwise comparisons identified by asterisks.

The number of observed adults was significantly related to time of day (LRT _3,99_ = 50.94, *p* = 0.001), the interaction between height and time of day (LRT _9,90_ = 25.15, *p* = 0.005, Fig. 6), and survey date (LRT _6,105_ = 30.86, *p* = 0.001). In the early morning, land counts and counts at 40 m assigned more adults than counts at 120 m (land, *p* = 0.049; 40 m, *p* = 0.033), but from late morning onwards, all surveys (land counts and drone counts at 40 m, 80 m, and 120 m) counted similar numbers of adults. The number of adults counted from the land and at 40 m did not significantly change with time of day, but for the other heights, fewer adults were counted in the early morning compared to late morning (120 m, *p* = 0.040), early afternoon (80 m, *p* = 0.004; 120 m, *p* = 0.003), and late afternoon (80 m, *p* = 0.016; 120 m, *p* = 0.003).

**Fig 6.**
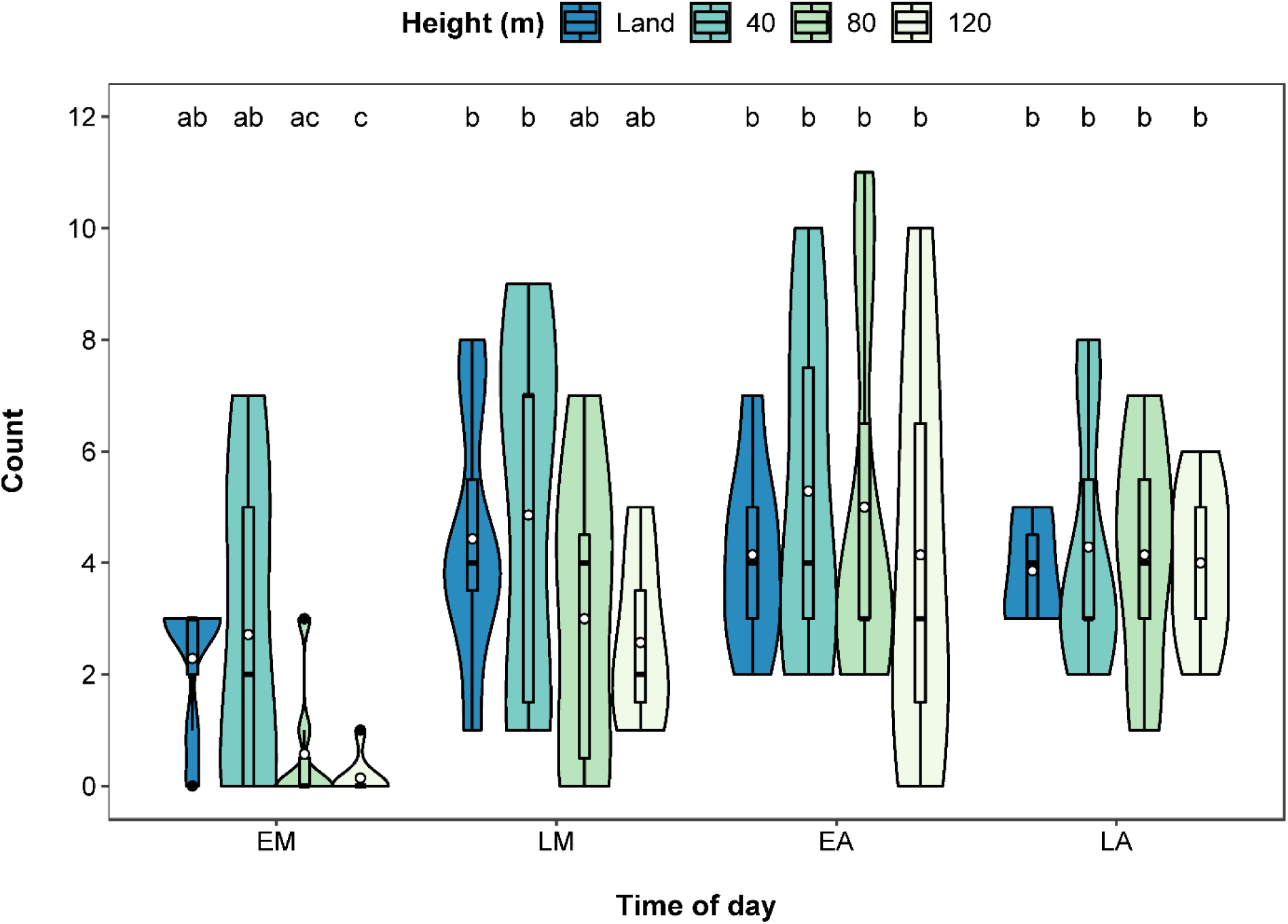
Violin and boxplot (mean, circle) showing significant interactive effect of height and time of day (EM – early morning, LM – late morning, EA – early afternoon, LA – late afternoon) on counts for adult hippos. Significant post hoc pairwise comparisons identified by letters.

### Discussion

We showed that by flying a relatively cheap drone at 40 m, as long as not in the early morning, reasonable estimates of hippo numbers and demographics were achievable, equal to or better than land counts. The effectiveness of a reasonably low height reflects the increased video resolution with decreasing height; at higher heights it is difficult to distinguish and count individual hippos (Fig. 2, S3 Figure). Land counts were possibly more accurate in our site than would be experienced in many other sites because the pod was reasonably habituated to humans and vehicles, given regular visits by tourists. Disturbance of less habituated hippos is likely in other places [58–60], leading to poorer land counts. Further, with increasing hippo pod size and water body size, land surveys become increasingly difficult, given uniformity of hippo appearance and diving behaviour [12].

Encouragingly, the drone did not disturb the hippos. We surveyed a relatively small pod (maximum 14 individuals), whereas hippo pods can sometimes number in their hundreds [46,61]. A larger pod size would still be relatively easy to survey using a drone, though it would take longer to count and differentiate demographic groups; time-consuming data processing is a drone cost [24,62]. Increasingly, such data processing could lend itself to automation through machine learning, which has already proven successful at identifying hippos on thermal infrared images [35], although it may be more difficult using RGB images, given the low colour contrast. Although hippos in larger congregations may be difficult to identify and track on video [36], our video continuously recorded the lagoon, allowing detection of hippos which surfaced momentarily, easily missed on images.

The lower height of the drone also allowed the demography of the hippo pod to be effectively estimated, with no significant difference in the percentage of hippos assigned to age classes between land counts and counts at 40 m (Fig. 4). We identified more juveniles and subadults from our land counts, probably because they were easier to see than from drone footage and were able to be visually assigned to age classes, even when they were partially submerged. This was reflected in the similarity between drone and land counts of adults, given their relatively larger size, and because we classified partially submerged hippos over a certain size as adults on drone images. In addition, the small sample size of juveniles (one) and subadults (two) may have reduced the statistical power of our analyses. Estimating demographic groups in the Democratic Republic of Congo pods had mixed success, with numbers of each age class varying for different flights over the same pod [34]. This could reflect differential hippo submergence between flights, unsuitable survey time of day, and visual extrapolation of body sizes. Restricting measurements to fully visible hippos reduces the sample size of hippos that can be assigned to age classes, but this can be increased by surveying when hippos are more exposed.

Aerial surveys of hippos are routinely flown at around 100 m at speeds of 160 - 180 km/hr [4,28,63–65], with observers estimating hippo numbers. It is unsurprising that these surveys underestimate hippos compared to land surveys [3,61,66]. There are clearly advantages to drone surveys; they capture hippo data at high resolution, given the relatively low flight height. The slower speed of the drone also increases viewing time, improving observations of hippos when diving and resurfacing. However, there are considerable advantages of the larger spatial coverage possible with aerial surveys, also accessing areas inaccessible for vehicles and drones. Inexpensive drones offer considerable promise for effective surveys of hippo pods, although battery life (about 20 minutes flight time) and flight range limits coverage to relatively small areas. The utility of drone technology is as an intermediate tool between lower accuracy, high cost, large scale aerial surveys and high accuracy but labour intensive land surveys [67].

Early morning was clearly an unsuitable time to effectively survey hippos (Figs. 2–6). This is when hippos were very active, continuously diving and surfacing, after returning to the water from nocturnal feeding [20,38]. This could also be a problem late in the day, when there is high activity [13,38,42], but was not detected because our drone surveys occurred before sunset. Our highest hippo counts were in the late morning and afternoon when hippos usually rested as a group during the middle of the day [20,38], often in shallow water with most of their body exposed, making them easy to detect and distinguish (S3 Figure). Our avoidance of early mornings for hippo drone surveys runs counter to recommendations from surveys of Democratic Republic of Congo hippo pods [36], although these did not test the effect of time of day. Instead, they argued for the advantages of minimising sun reflection, which we effectively reduced by recording video, and surveying when they considered hippos most visible [27]. Hippo behaviour may differ by region or habitat, we therefore recommend adapting the timing of surveys to when hippo are resting, which may vary in location and time. Importantly, our surveys also effectively tracked changes in the hippo pod over time, as adults emigrated from the lagoon as it dried, a typical response of hippos to changing water availability [5,26].

Drones are increasingly valuable for monitoring wildlife populations [24,33,68], including hippos. Our analyses show that drone data can provide accurate estimates of hippo pods, including their demographic structure. Importantly, they also provides a viable alternative to land based counts, with low impact on hippos, offering further opportunities to survey in difficult to access areas [3,26,61] and, just as critically, collect these data safely. Such data could be routinely collected in different river systems, providing indices of abundances, temporal changes and tracking the long-term status of hippo populations, an imperative given their declining populations in many parts of Africa.

## Acknowledgements

We thank Elephants Without Borders for hosting the study and providing logistical support. In addition, we thank the Botswana Ministry of Environment, Wildlife and Tourism for affording us the opportunity to conduct this research. The University of New South Wales and the Australian Government supported our work. We thank Fly UAS for sponsoring Remote Pilot Licence training, Keboditse “CK” Mboroma for his assistance in the field, and Wilderness Safaris and staff at Abu and Seba Camps.

## Supporting information

**S1 Table. Complete dataset.**

**S2 Text. Results of statistical models, as outputs from R.**

**S3 Figure. Snapshots of survey videos at a) 40 metres, b) 80 meters, c) 120 metres.** Notice the increasing difficulty of detecting hippos with increasing altitude due to lowering resolution. Images taken during early afternoon surveys showing resting posture of hippos with the majority of their body exposed, allowing easy detection and counting.

